# Stochasticity of cellular growth: sources, propagation and consequences

**DOI:** 10.1101/267658

**Authors:** Philipp Thomas, Guillaume Terradot, Vincent Danos, Andrea Y. Weiße

## Abstract

Cellular growth impacts a range of phenotypic responses. Identifying the sources of fluctuations in growth and how they propagate across the cellular machinery can unravel mechanisms that underpin cell decisions. We present a stochastic cell model linking gene expression, metabolism and replication to predict growth dynamics in single bacterial cells. In addition to several population-averaged data, the model quantitatively recovers how growth fluctuations in single cells change across nutrient conditions. We develop a framework to analyse stochastic chemical reactions coupled with cell divisions and use it to identify sources of growth heterogeneity. By visualising cross-correlations we then determine how such initial fluctuations propagate to growth rate and affect other cell processes. We further study antibiotic responses and find that complex drug-nutrient interactions can both enhance and suppress heterogeneity. Our results provide a predictive framework to integrate single-cell and bulk data and draw testable predictions with implications for antibiotic tolerance, evolutionary biology and synthetic biology.

## I. INTRODUCTION

The rate at which cells accumulate mass and grow is highly variable across isogenic cells^1–4^. Previous studies have considered fluctuations in growth rate as one of the major drivers of phenotypic heterogeneity^4–7^. Yet the physiological origins of these fluctuations remain elusive so far. Growth laws characterise the typical behaviour of cell populations^8^, for example, the scaling of average growth rate with macromolecular composition, including ribosome and other protein levels, or cell mass in bacteria^8–10^. These phenomenological relations can give important insights into the population average behaviour, but may not translate to an understanding of individual cell responses^3^.

There is substantial evidence that cellular noise sources are diverse and may propagate via growth in a systemic way. A recent experimental study showed that fluctuations in the expression of enzymes caused considerable variation in the growth rate of single cells, which then fed back onto their expression and that of other genes^1^. Cell-to-cell differences stem from intrinsic fluctuations in biochemical reactions^11^. Some of these reactions, particularly those that drive cell growth, affect many other intracellular processes, and so a range of cellular responses can vary even under constant conditions^12^. Gene expression, for example, is known to be inherently stochastic at the single-cell level^11^. It is less clear though how variations affect regulatory mechanisms that control intracellular processes^13,14^, and how this translates to phenotypic differences and cell fitness.

Models can help us identify potential sources of growth fluctuations and understand how they propagate to cause phenotypic variation. There are various approaches to model cellular growth. One is to invoke growth rate optimisation^15–17^, others consider the coordination of growth with gene expression^8^ or combine the two approaches^18^. Such approaches have been used to model static cell-to-cell variation by imposing parameter variability onto the model behaviour^19,20^. The sources of growth variations, however, remain unclear, and also how to adapt the models to explain cell responses that fluctuate over time.

Here we present a stochastic model of single-cell bacterial dynamics to predict the growth rate of individual cells. Our description of cells is based on biochemical kinetics, which can more amenably account for how heterogeneous responses arise from stochastic fluctuations in cellular mechanisms. In this context, the magnitude of fluctuations results from the abundance of key molecular players^21^, and so we can avoid assumptions of variability imposed onto the model behaviour.

The model builds upon recent mechanistic insights into population-average growth responses via a coarse-grained description that explains Monod-growth and empirical relations between growth rate and ribosomal contents from the interplay of nutrient uptake, metabolism and gene expression^8,22^. These processes are subject to constraints by cellular trade-offs such as a finite transcriptome and proteome per cell as well as limited pools of ribosomes and cellular resources. Here we consider the finite number of intracellular molecules produced over a cell-cycle and so explicitly account for the accumulation of cell mass and its corresponding stochastic dynamics. We further integrate this approach with a simple model of bacterial cell-cycle control^23,24^, as supported by recent experiments^4,25^, and quantitatively predict emergent growth and division dynamics in single cells.

Along with the cell model we present a theoretical framework to approximate stochastic growth and division dynamics based on a small noise approximation. The framework is general, that is, applicable to models of reaction-division systems at large. It enables closed-form computation of model statistics, such as mean and variance of variables over time, and thus allows efficient parameter estimation from single-cell data alongside a systematic decomposition of the sources of growth variation.

Our modelling approach, in combination with the developed approximation, allows us to statistically characterise the macromolecular composition, growth rate and mass of single cells. It recovers several empirical responses at the population- and single-cell level, thus providing substantial validation. We quantify the contributions of different noise sources to observed growth rate fluctuations and analyse their propagation. As a result, we identify noisy dynamics of mRNAs coding for nutrient transporters and enzymes as a major source of growth rate fluctuations. We moreover find that growth rate is a source of global noise that can be transmitted to other processes, for example, via ribosomes^6,26,27^. Our analysis of cell responses to translation-inhibiting antibiotics further indicates a strikingly complex dependence of growth heterogeneity on environmental conditions, which may pinpoint strategies to avoid drug tolerance.

## II. RESULTS

### A. Quantifying single-cell growth

#### A stochastic model of single-cell growth

Developing models that coordinate growth and division in single cells is a major challenge because they need to integrate many processes at different scales. We take a hybrid approach to model growth and division of single cells by integrating DNA-replication with stochastic biomass production (Fig. 1a).

Cellular protein content dominates biomass, and thus the total translation rate determines the rate of biomass production^8,22^. In a single cell, translation is coupled to processes that fuel and drive gene expression. Since these processes are stochastic, growth rates varies over time and from cell to cell. We use a bottom-up approach that describes the dynamics of a coarse-grained cellular composition based on stochastic biochemical reactions, which comprise transcription, translation, ribosome binding, mRNA degradation and metabolism (see Methods). The model describes the accumulation of proteomic mass–split into sectors containing transporters (*t*), catabolic enzymes (*e*), ribosomes (*r*) and housekeeping proteins (*q*)–along with the corresponding transcriptome, ribosome-mRNA complexes and a resource molecule. The resource is a coarse-grained variable describing the collection of molecules that fuel biosynthesis, for example, energetic molecules such as ATP and NAD(P)H or charged tRNAs, depending on the nutrient limitations under consideration. It can then be shown that the kinetics recover Monod-growth, that is, growth rate saturates for increasing nutrient qualities^8,22^.

**FIG. 1.**
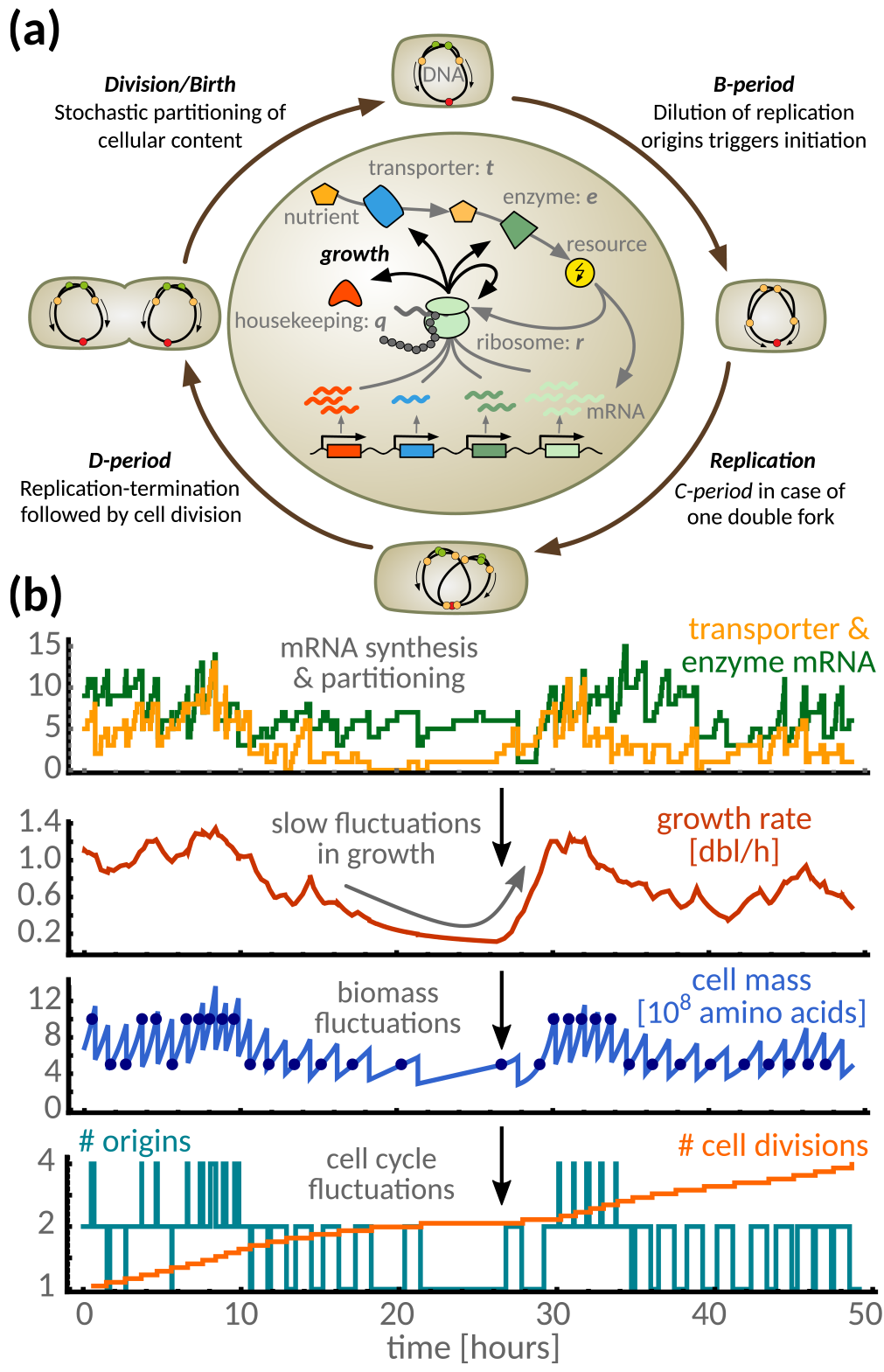
Stochastic model of single-cell growth. **(a)**The outer cycle illustrates the cell cycle model based on the Cooper-Helmstetter model of bacterial replication. We assume initiation of a new round of replication at a fixed concentration of DNA-origins, analogous to a fixed initiation mass per DNA-origin^24^, thus growth dynamics schedule the replication events and are determined by the intracellular model (inner circle). The latter describes import and metabolism of resources, and how they fuel gene expression, where the rate of protein-biosynthesis determines growth. Stochasticity of cellular dynamics is a result of the intrinsic stochasticity of the various reactions and the random partitioning of the cellular content at division. **(b)** Stochastic simulations illustrate the propagation of intrinsic fluctuations in single cells: mRNAs are synthesised at low numbers per cell (yellow & green lines), which affects protein production and so growth rate (red line). Fluctuations in growth lead to temporal variations in cell mass that can span several cell cycles (blue line), causing fluctuations in the number of replication origins (teal line), in the mass at initiation (filled circles), and consequently in cell divisions (orange line).

At the single-cell level, we must also account for cell divisions. We follow the Cooper-Helmstetter model in which cells divide after a constant period following initiation of DNA replication^23^. Allowing for parallel replication rounds, replication cycles couple to growth through initiation at a fixed concentration of replication origins^24^. As a consequence, the cell mass at initiation depends on the number of ongoing replication rounds in a given growth condition (Methods b).

To capture the stochastic dynamics of the model we focus on a lineage description that tracks a single cell over many replication and division cycles. At division, all intracellular molecules are partitioned randomly between the two daughter cells, and we retain information about only one daughter cell^28^. We account for asymmetric cell divisions, as for instance due to inaccurate positioning of the division septum^2,3,29^, and assume that molecules are partitioned according to the inherited volume fraction of the daughter cell (SI Sec. I).

Fig. 1b illustrates the dynamic propagation of fluctuations using stochastic simulations: The stochastic synthesis, degradation and partitioning of mRNA molecules lead to slow fluctuations in the growth rate and in turn to variations in biomass production and interdivision timings. Simulations of this stochastic model are computationally expensive due to the large number of molecules produced per division cycle. We therefore developed an approximation method that facilitates quantitative insights into a whole class of cell models involving coupled reactions and divisions.

#### Stochastic analysis of reaction-division systems

Consider a generic reaction-division system composed of *N* intracellular species with molecule numbers *x* = (*x*_1_, ‥, *x*_*N*_). The macromolecular composition of a single cell determines its mass via

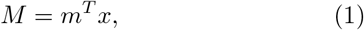

where the components of the vector *m* denote the mass of individual molecules. At constant macromolecular density this measure is directly related to cell size. The corresponding intracellular concentrations are given by

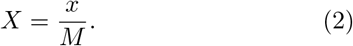

For an intracellular reaction network comprising *R* reactions with stoichiometric matrix *ν*, the cell growth rate can be obtained analytically and is given by

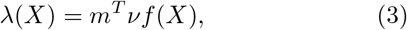

where *f*(*X*) is the vector of reaction rate functions (see SI Sec. II). Because the intracellular reactions are stochastic, the concentrations *X* fluctuate over time from which follows that the growth rate is a stochastic process.

We characterise the dynamics of intracellular concentrations, cell mass and its growth rate using a continuous approximation that describes biochemical reactions, dilution due to overall biomass production, and partitioning of molecules at cell division. The set of coupled Langevin equations is

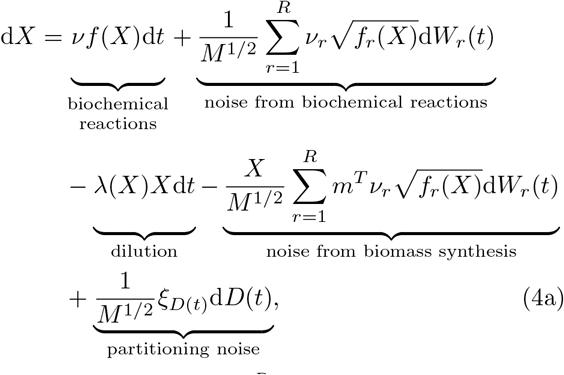

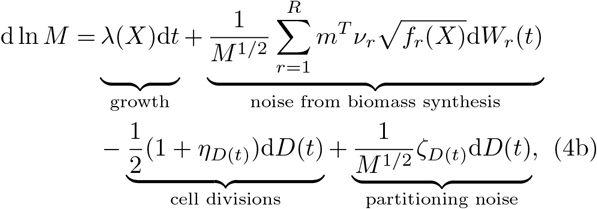

which involves a process *D*(*t*) counting the number of divisions (see Fig. 1b, SI Sec. I), independent Gaussian white noises *W*_*r*_(*t*) describing the intrinsic variability of the intracellular reactions, the random variables *ξ*_*D*_ and *ζ*_*D*_ introduced from partitioning of molecules at cell division, and *η*_*D*_ from the variation in the inherited volume fraction (see SI Sec. IIA for a detailed derivation). Specifically, for every cell division the noise terms satisfy 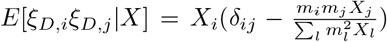, 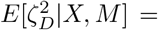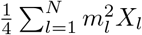 and 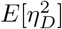 equals the squared coefficient of variation of the inherited volume fraction. In the deterministic limit, that is, for large *M*, the concentration process, Eq. (4a), is independent of cell mass. In the stochastic case, however, the two processes are coupled.

In between cell divisions (d*D* = 0), Eq. (4b) gives the instantaneous cell growth rate, which has two contributions: λ(*X*), a function of intracellular concentrations, and a second random part that stems from the mass-producing reactions. We find that for biologically relevant situations the second contribution is negligible due to averaging over the large number of such reactions occurring between cell divisions (SI Sec. IIA). We note that, in the absence of growth, that is, when all reactions are mass-conserving (*m*^*T*^*ν*_*r*_ = 0), Eqs. (4) reduce to the standard chemical Langevin equation^31^.

To gain further analytical insights we developed a small noise approximation^32,33^ of Eqs. (4). The approximation allows us to obtain mean concentrations and growth rate by solving a coupled system of ODEs in steady-state conditions. Concentration fluctuations lead to growth rate variations that can be computed in closed form (Methods b). The method provides accurate estimates of the first two statistical moments (Fig. 2), and thus enables efficient inference of model parameters, which is typically infeasible using stochastic simulations^34^. We discuss the results and predictions drawn from this inference using experimental bulk and single-cell data (Methods c) in the following.

**FIG. 2.**
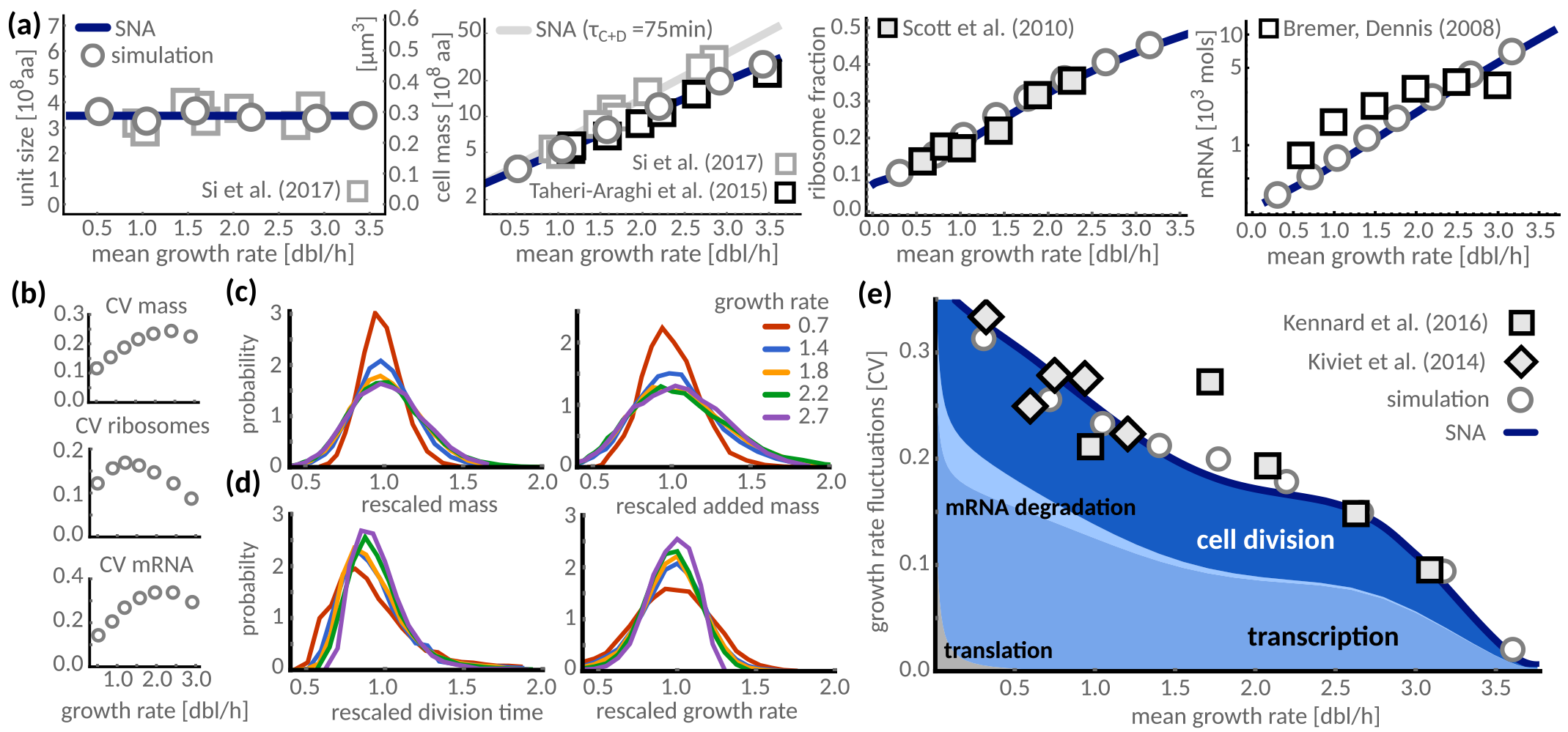
The mechanistic model predicts condition-dependence of growth rate fluctuations in single cells. **(a)** The stochastic model recovers common bacterial growth laws. Cell mass per number of origins (unit size) is constant in all growth conditions. Absolute cell mass, ribosome content and total mRNA numbers per cell increase with average growth rates. Filled markers denote measured quantities used to parametrise the model (see SI for details), empty markers denote measurements that validate model predictions. Circles are the results of stochastic simulations and validate the employed approximations (SNA). A longer C+D-period of 75 minutes as observed in^30^ yields higher predictions for cell mass (grey line). **(b)** Fluctuations, measured by the coefficient of variation (CV), in cell mass initially increase as a function of average growth rate. Fluctuations of ribosomal mass fraction are of the order of 10 – 20%, and those of total mRNA concentrations largely follow the trend of the mass CV. **(c)** Single-cell distributions of cell mass at birth and mass added between birth and division are invariant when rescaled by their means. For intermediate to fast growth conditions (1.4-2.7 doublings per hour) distributions collapse nearly perfectly, consistent with the stable CV in this growth regime (b), while slowly growing cells (0.7 red line) deviate from this universal behaviour. **(d)** In contrast, the distributions of rescaled doubling times and growth rates broaden gradually with decreasing medium quality, highlighting that these quantities are condition-dependent at the single-cell level. **(e)** Our model quantitatively recovers variations over the whole range of experimentally accessible growth rates. In agreement with experimental observations (squares^3^, diamonds^1^) fast growing cells display less growth variability than slow growing cells. This dependence is well predicted by stochastic simulations (grey circles) and by the small noise approximation (SNA, solid blue line). Colours indicate the contributions of different cellular processes to growth variations: synthesis, degradation and random partitioning of mRNAs at cell division. The contributions from other processes such as protein translation are overall small (grey area).

### B. Condition-dependence of growth in single cells

#### Macroscopic growth laws

The macromolecular composition of *E. coli* is growth-rate dependent, and we ask whether the cell model is consistent with several bacterial growth laws describing these relations. Our model predicts that mean cell mass increases exponentially with mean growth rate, the Schaechter-Maaløe-Kjeldgaard growth law^9^, as a consequence of the coupling of DNA-replication to growth^24^. We moreover find that unit size^30^, in terms of mass per number of origins, is invariant across growth conditions (Methods b, Eq. 8). If we compare the theoretical unit size with recent measurements of unit volumes in *E. coli*^30^ (Fig. 2a, first panel), the model predicts a protein density of 12×10^8^aa/*μm*^3^, well in line with literature values^35^. With this density estimate, model predictions closely match cell sizes reported in two different datasets^2,30^(Fig. 2a, second panel). The inferred model further recovers ribosome abundances over the experimental range of growth rates^8^ (third panel) and predicts transcriptome and proteome sizes that are in qualitative agreement with experimentally observed values^36,37^ (fourth panel, and SI Fig. S3b).

**FIG. 3.**
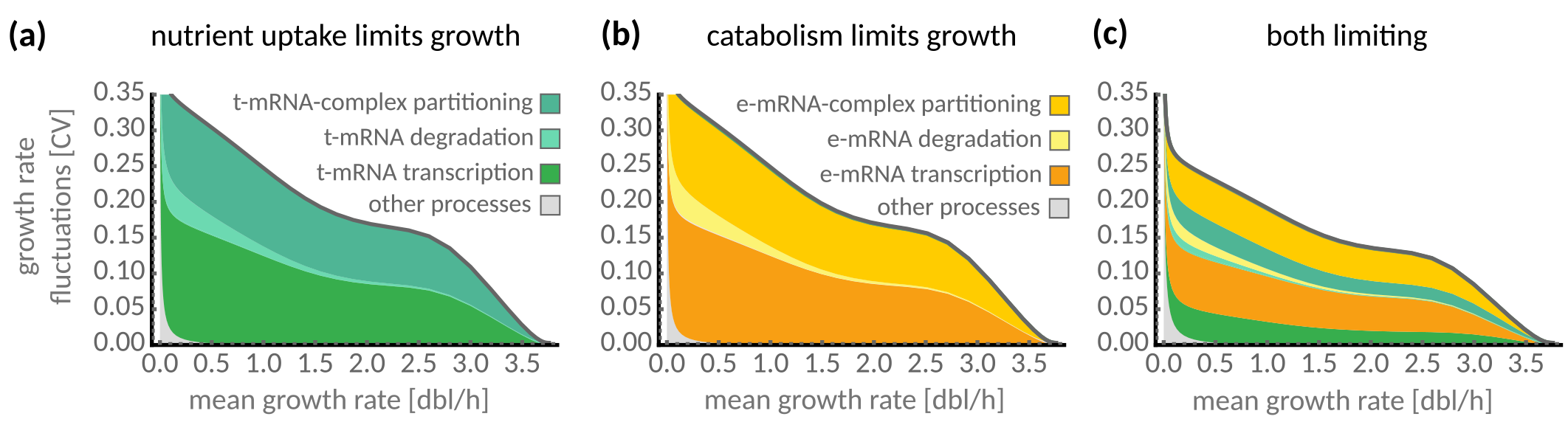
Nutrient uptake and catabolic limitations determine the sources of growth fluctuations. We vary catabolic turnover rates *υ*_*m*_ relative to nutrient uptake rate *υ*_*t*_. **(a)** Fluctuations in transcription of transporter mRNAs and their stochastic partitioning dominate growth variability because nutrient uptake is growth-limiting (*υ*_*t*_ > *υ*_*m*_). Other processes such as translation of proteins are largely negligible. **(b)** When catabolism limits growth (*υ*_*m*_ > *υ*_*t*_), fluctuations in enzymatic mRNAs instead explain most of the observed variation. **(c)** For comparable uptake and turnover rates (*υ*_*m*_ ≈ *υ*_*t*_), both transporter and enzyme mRNA fluctuations contribute to the overall variation. The absolute size of fluctuations reduces indicating a noises cancellation effect.

#### Scaling of single-cell distributions

Recent experiments suggest a universal behaviour of cell size and doubling time distributions when rescaled by their mean^2,3^, indicating that growth conditions primarily affect the mean cell size, doubling time, and growth rate. Our model reproduces this dependence for the cell size and added mass in intermediate to fast growth conditions (Fig. 2b,c). For conditions slower than those measured in^3^ we observe a breakdown of this scaling. In those conditions, our model predicts an increase in cell size variability with growth rate (Fig. 2b) due to a shift from single to parallel rounds of replication^4,38^(SI Fig. S3a). Such increases in variations of cell mass have indeed been observed in cells grown in the mother machine^2,38^. Similarly, we observe no scale invariance for doubling times and growth rates (Fig. 2d), indicating that their cell-to-cell variations are condition-dependent rather than universal. The model in fact explains an empirical condition-dependence as discussed in the following.

#### Condition-dependence of growth rate fluctuations

Recent data suggest a condition-dependence of growth rate fluctuations in single *E. coli* cells^1,3^ (Fig. 2e). In line with these observations, our model predicts growth variations to decrease with mean growth rate. This dependence is well captured by the developed approximations and stochastic simulations (Methods d). Our model predicts that fluctuations vanish as the mean growth rate approaches its maximum. This is not because intracellular reactions stop fluctuating, but rather because growth rate saturates and thus such fluctuations no longer translate to growth variability.

### C. Sources of growth rate fluctuations

#### Processes contributing to growth fluctuations

Our model allows us to investigate the sources of phenotypic variations. We use a noise decomposition^39^ to study the effect of each reaction and partitioning at division on the overall noise levels (Methods b). We find that transcription and cell division are the major determinants of growth heterogeneity across all growth conditions (Fig. 2e). Degradation of mRNA only becomes important at slow growth, when its rate dominates over dilution. Nutrient uptake and metabolism, in turn, yield negligible contributions because nutrients are highly abundant (SI Fig. S6). Similarly, effects of noise in translation are mostly negligible, due to the large number of proteins synthesised during a cell-cycle, and only contribute to growth variations at very small growth rates. In such conditions, however, regulatory mechanisms as involved in starvation are expected to take effect which our model does not describe.

#### Limiting factors to growth

The dominant contribution to growth variations stems from the synthesis and removal of transporter mRNAs (Fig. 3a). This suggests that nutrient uptake limits growth rate, consistent with estimated catabolic rates exceeding those of nutrient uptake (SI Tab. S1). Since catabolic rates can be tuned by cofactors^41^, we wondered whether limiting catabolic turnover could affect growth fluctuations. When catabolic turnover is slower than nutrient uptake, indeed, growth variations are due to the transcription and removal of enzyme mRNAs rather than transporter mRNAs (Fig. 3b). The total size of growth fluctuations remains largely unaffected by whether uptake or catabolism limits growth. Surprisingly though, when both nutrient uptake and catabolic turnover are simultaneously rate limiting, growth variability dips, suggesting a noise cancellation effect (Fig. 3c, SI Fig. S4b). Operon organisation of the corresponding genes does not affect our predictions, except that the simultaneous limitation by transport and catabolism does not lead to noise cancellation (SI Fig. S4).

**FIG. 4.**
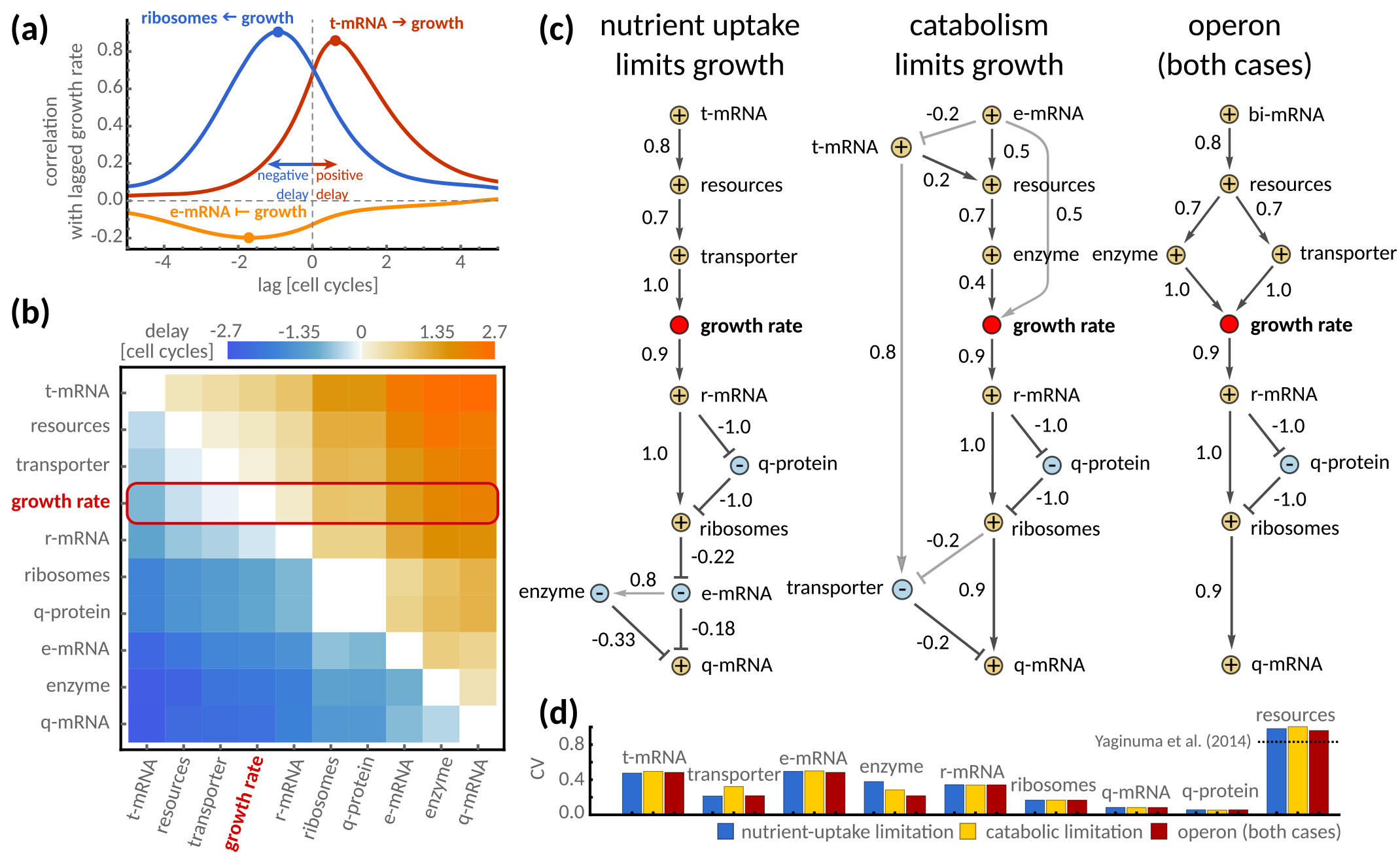
Cross-correlation analysis reveals the propagation of fluctuations. **(a)** Fluctuations in the concentrations of transporter mRNA (t-mRNA) correlate positively with growth rate at later times (red line, maximum correlation at positive lag) indicating that they increase growth rate. Ribosome concentrations correlate positively with growth at earlier times (blue line, maximum correlation at negative lag) indicating that they are increased by growth rate. Concentrations of enzymatic mRNA (e-mRNA) correlate negatively, indicating that they are mainly diluted by growth (yellow line, minimum correlation at negative lag). **(b)** Pairwise delays between cellular components and growth rate (red box), ordered by their delay with respect to growth rate and computed in moderate growth conditions. **(c)** ‘Minimal delay graphs’ illustrate the information flow under different growth limitations with comparable growth rates. A dark arrow from species A to B indicates that B has minimal delay from A, and so species B are the first to receive fluctuations from species A. Arrows denote positive correlations at the delay, T-arrows negative correlations, and the label denotes the delayed-correlation coefficient. We include light arrows to indicate components with second smallest delay whenever these are not reached through subsequent steps. The graph reveals cellular components up-and downstream of growth rate, i.e. those that affect growth and those affected by growth. When nutrient-uptake is limiting growth, t-mRNA act as a source of fluctuations, while for catabolic limitation e-mRNA are the dominant source (cf. Fig. 3). Their corresponding proteins are upstream of growth and transmit fluctuations to growth rate under the respective limitations. When transporters and enzymes are co-expressed from an operon these limitations are indistinguishable. In all cases, q-mRNAs act as a sink due to their negative auto-regulation, q-proteins are mainly diluted (nodes labelled with – correlate negatively with growth) while most species increase with growth (nodes labelled with + correlate positively with growth). **(d)** The size of fluctuations (CV) in concentrations of the transcriptome, proteome and resources is comparable in the above cases. The dashed line indicates measured fluctuations in intracellular ATP^40^.

Having identified metabolic components as limiting factors to growth and as a major source of variation, it is important to consider their absolute abundances, because they determine the magnitude of noise^12^. In our model, mean abundances of transporter and enzyme mRNAs vary in different conditions between 3 and 9 copies per cell, with approximately 9, 000 molecules of the corresponding proteins (SI Fig. S6). Compared to that, natural abundances are between 10^−3^ and 1 mRNA copies per gene while proteins are more abundant with 1 to 10^3^ molecules, with products of essential genes occurring at higher numbers^42^. This suggests that transporter and enzyme species in our model are consistent with lumped groups of molecules rather than a single rate-limiting species.

### D. Propagation of fluctuations

We further ask how stochastic fluctuations propagate to growth, and how this affects the macromolecular composition of a single cell. Since all intracellular concentrations are interconnected and growth rate feeds back onto them via dilution, it is not straightforward to determine the flow of information. We use cross-correlation across the different intracellular species and growth rate, computed from stochastic simulations, to quantify the propagation of fluctuations. Cross-correlation measures the similarity of two quantities at different time instances. The time at which maximal correlation is reached measures the delay between correlations and marks their temporal order, and so cross-correlation can indicate causality^1^. The model does not comprise effects that are detrimental to growth. Any upstream components, that is, species transmitting fluctuations to growth, are therefore expected to promote growth and so correlate positively with it. Growth in turn can either increase or dilute downstream components, and so its correlation with downstream components may be positive or negative.

**FIG. 5.**
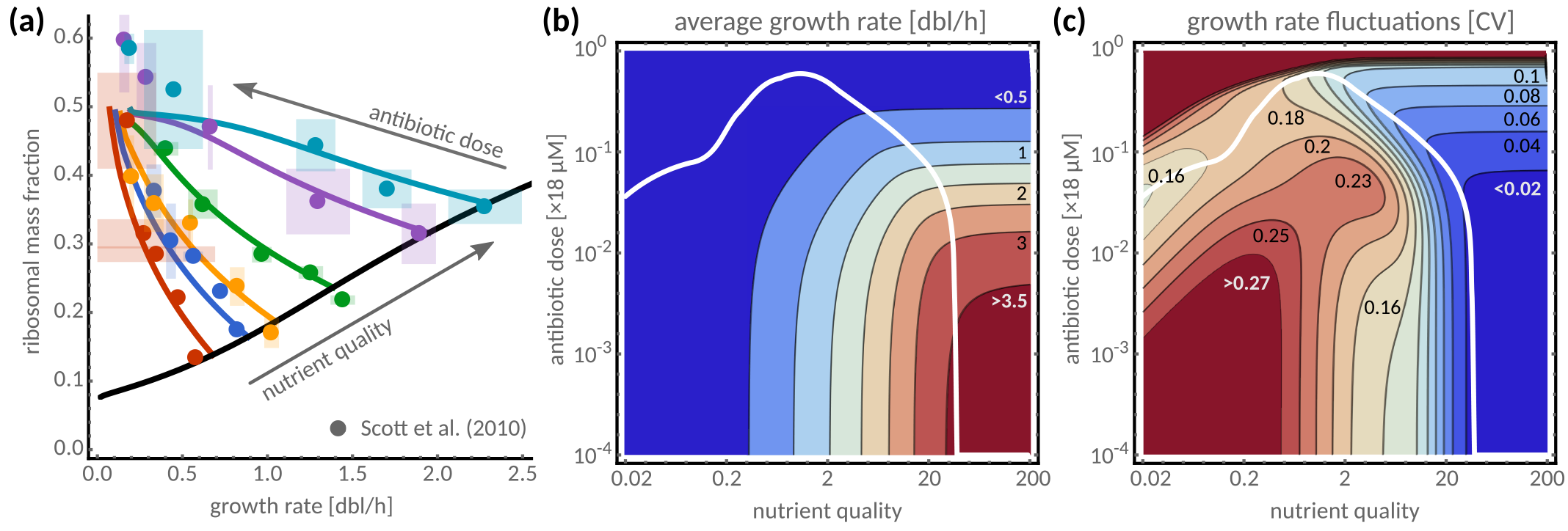
Condition-dependence of antibiotic responses. **(a)** Ribosomal content per cell as a function of average growth rate after treatment with chloramphenicol. Predictions (solid lines) are in quantitative agreement with experimental data^8^ (dots, shaded areas denote standard deviations over replicates and colours denote different nutrient conditions). **(b)** For any given nutrient condition the average growth rate is predicted to decrease monotonically with antibiotic dose. **(c)** Growth heterogeneity is predicted to be highly complex in both a nutrient-and dose-dependent manner. In nutrient-rich conditions growth heterogeneity increases with antibiotic dose, while in intermediate and poor conditions the response is non-monotonic. Over a large range of nutrient conditions there exists a non-zero drug dose that minimises growth heterogeneity (solid white line, b and c).

We observe a strong positive correlation of transporter mRNA concentrations with growth rate at later times (Fig. 4a), consistent with our previous finding that their fluctuations are the major source of growth variations. Ribosome concentrations also correlate positively with growth, consistent with the increase of mean levels with growth rate (compare Fig. 2a). Interestingly they correlate at a negative delay, suggesting that fluctuations in growth propagate to ribosomes but ribosome fluctuations do not contribute substantially to growth variability. Other species such as enzyme mRNAs correlate negatively at a negative delay, indicating their concentrations are mainly affected by dilution, a relation that we observe more generally for their corresponding enzymes and also for q-mRNA across all conditions.

To estimate the propagation of fluctuations in the upstream and downstream processes of growth we consider the delay between any pair of groups (Fig. 4b). The intuition behind this is that a minimal positive delay indicates the species that first ‘senses’ a fluctuation, which it then passes on to the next species.We illustrate this flow of information in a directed graph, where edges indicate the minimal delay relation between groups of species and edge weights their correlation (Fig. 4c).

Consistent with the noise sources identified in Fig. 3 we find that, depending on whether nutrient uptake or catabolism is growth-limiting, fluctuations in transporter or enzyme mRNAs are the source of growth rate variation that propagate via their respective protein levels and resources to growth. Further, when transporters and enzymes are co-expressed from an operon, as in^1^, their common mRNA is the dominant source of growth rate variations and the noise propagation in the two limiting regimes is indistinguishable. All other components are downstream of growth rate, steadily across different growth conditions (SI Fig. S5a), and are thus affected by growth. Only at high growth rates ribosomal transcripts - but not their proteins – are upstream of growth, because in these conditions fluctuations in ribosomes rather than in resources dominate noise in growth rate (SI Fig. S5, cf. Eq. (6)). Interestingly, q-mRNAs act as noise sinks as they are subject to negative auto-regulatory control.

Highly abundant species have consistently lower noise levels (Fig. 4d, SI Fig. S6) except resources, which display an extremely high variability, likely due to their central role in many cellular reactions. This prediction is quantitatively confirmed by recent experiments showing that ATP-levels in *E. coli* vary up to 80%^40^ (compare Fig. 4d). Our model suggests that such large fluctuations can impact growth, as has been observed in eukaryotes^43^. The analysis shows that growth rate affects a large number of downstream components, which may include, for example, transcription factors controlling stress responses or other phenotypic switches. Our results therefore suggest that growth rate plays a central role as a source of global noise driving phenotypic heterogeneity.

### E. Growth heterogeneity in response to antibiotics

To showcase the utility of our approach we next examine bacterial responses to antibiotic treatment. The common route to assess the efficacy of drugs is by establishing the dose-dependence of growth rate^44^. Growth heterogeneity, however, is also important as it gives rise to antibiotic tolerance, which allows individual cells to survive treatment through non-genetic mechanisms^45–47^. Surviving cells then have sufficient time to develop and pass on mutations that confer resistance to later treatment, and so growth heterogeneity significantly contributes to the rise of antibiotic resistance. Ideally, treatment should therefore aim to minimise pathogenic growth while avoiding regimes of high growth heterogeneity.

Our model correctly predicts average macroscopic composition in various nutrient conditions and in response to the ribosome-targeting antibiotic chloramphenicol (Fig. 5a). We thus use it to quantitatively map both average growth rate and growth heterogeneity to combinations of nutrient and antibiotic regimes. We modelled the action of chloramphenicol via inactivation of ribosome complexes (Methods a). Without re-fitting, the model predicts that average growth rate increases with nutrient quality and decreases with antibiotic dose (Fig. 5b).

Predictions for growth heterogeneity display a more complex response (Fig. 5c). For all nutrient conditions growth heterogeneity rises steeply at high drug concentrations. But only in very rich nutrient conditions, where growth rate saturates, growth heterogeneity increases monotonically with antibiotic dose, consistent with our previous predictions that average growth and heterogeneity exhibit an inverse relation (cf. Fig. 2d). In all other nutrient conditions, growth heterogeneity is highly non-monotonic as a function of dosage: In medium-to-rich nutrient conditions, heterogeneity first peaks and then dips before the final rise. In low-to-medium nutrient conditions the final rise is preceded by a drop in growth heterogeneity at intermediate doses.

Our predictions suggest that avoiding regimes of high growth heterogeneity may be achieved in different ways depending on the location of an infection. For example, it may be possible to treat infections in low-to-medium nutrient conditions, such as the urinary tract or blood, with a dose that minimises heterogeneity (Fig. 5c, white line). This would require more care for infections of richer nutrient environments, such as the gut, where regimes of increased heterogeneity should be avoided. The predictions further suggest that infections of very rich environments cannot be treated with an overall reduction of growth heterogeneity. Notably though, heterogeneity is mostly low in these conditions, and so an ideal dose may be chosen high enough to sufficiently inhibit growth but low enough to avoid regimes of significant heterogeneity. Alternatively, treatment efficacy may be manipulated by changing the environment of the pathogen, for example, by constraining diet.

## III. DISCUSSION

We presented a stochastic cell model to predict growth and division dynamics in single bacterial cells. Our model yields detailed predictions of measurable macroscopic quantities including growth rate, size and macromolecular composition. In contrast to previous approaches, we predict phenotypic heterogeneity as it arises from the intrinsic fluctuations of biochemical reactions, and therefore as an emergent physiological response of single cells. Our approach is one of the first to predict cellular physiology and heterogeneity from molecular mechanisms and in response to complex environments that include different nutrient conditions and drug doses.

We quantitatively recover levels of growth heterogeneity that have been measured in individual bacterial cells, and predictions are in good agreement with absolute proteome, transcriptome and ribosomal levels per cell as reported in bulk measurements. We observe scale invariance of several macroscopic quantities over the range of experimentally reported conditions, indicating that cell-to-cell variations are independent of these growth media. Our results moreover suggest that this scale invariance breaks down if tested over a broader range of growth conditions. In particular, we predict an increase of cell size variability with growth rate, in agreement with recent experiments^2,38^.

We presented a general framework to analyse stochastic reaction-division dynamics. Our theoretical analysis enabled us to dissect the contributions of different biochemical processes to the observed growth heterogeneity. Specifically, we identified fluctuations in the synthesis, partitioning and degradation of mRNAs coding for proteins involved in metabolism as the major source of growth heterogeneity. The prediction is in line with observations that mRNAs of essential genes can naturally be present at low molecule numbers per cell^42^ and that fluctuations in enzyme expression can cause growth rate variation^1^. In fact, expression of glucose transporters in *E.coli* has been reported to be highly heterogeneous^48^.

In agreement with experiments we find that overall growth variability is condition-dependent, decreasing generally with mean growth rate. However, for mixed environments that involve nutrient-drug interactions we predict complex responses, where antibiotic doses can either decrease or increase variability depending on nutrient conditions. Our analysis moreover showed that different limitations to growth, including nutrient uptake and catabolism, can result in the same growth phenotypes, underlining the robustness of the predicted behaviour.

We analysed the propagation of stochastic fluctuations. We used a novel strategy to distinguish components upstream or downstream from growth, that is, cellular components transmitting fluctuations to growth or receiving them from growth, by building minimal delay graphs from pairwise cross-correlations. Species upstream of growth, such as transporter mRNAs and proteins as well as resources, correlate positively with growth rate, whereas species downstream of growth either increase with growth or are diluted. These predictions on the propagation of stochastic fluctuations may be tested using protocols similar to those employed by Kiviet *et al.*^1^.

In particular, we identified resources to exhibit significant fluctuations, which they transmit to growth rate. In support of this, recent experiments showed that intracellular ATP levels can indeed vary substantially^40^, and such variations can affect growth rate in eukaryotic cells^43^. We moreover found that ribosome levels correlate positively with growth rate, consistent with the known growth law^8^. But our model finds that ribosome fluctuations follow those of growth rate, in agreement with the observation that asymmetric ribosome partitioning at cell division has negligible effect on growth rate^4^. Our finding that ribosome levels are set by growth rate at the single-cell level moreover suggests that growth fluctuations are a common source of cellular noise, and ribosomes act as transmitters of this noise to downstream processes^6,26,27^.

The developed single-cell model also allowed us to identify biological parameters that are otherwise non-identifiable using deterministic population-averaged approaches^34^. For example, total resource levels, as opposed to concentrations, have little to no effect on mean growth but affect growth rate variances, which can only be constrained by single-cell data (SI Fig. S2). Recent work showed that such fluctuations can impact the mean population growth^49^, implying that these mechanistic parameters may be subject to evolutionary pressure. Cellular physiology could seize this degree of freedom and use it to shape noise to its benefit, for instance, as an evolutionary bet-hedging strategy^50–52^. The model could be put to use in *in numero* evolutionary experiments to test the specific benefits of noise architectures in different environments and retrace possible evolutionary paths.

Our framework may also prove useful to benchmark the designs of synthetic circuits and increase their reliability. In this context one may, for example, wish to limit the impact of growth fluctuation on a circuit of interest^53^. Embedding such circuit in our model provides insights to re-architecture the global cellular noise to this effect. Finally, our results have important implications for drug tolerance and could pinpoint strategies to potentiate clinical treatment. Increased cell-to-cell variability can also drive phenotype switching. The latter plays a crucial role in persistence, a form of tolerance that allows bacteria to survive antibiotic treatment by switching to a dormant state^46^, the precise mechanisms of which, however, are still unclear.

We limited our analysis to the effect of intrinsic fluctuations in the biochemical processes that underpin growth and neglected potential variations in processes responsible for DNA-replication and cell cycle control^4^. We further focussed on the balance between catabolic and biosynthetic processes^8,54^, where we considered effective regulation through the dependence of transcription on cellular resources (Methods a, see also^22^) and avoided mechanistic detail such as regulation via (p)ppGpp. In this sense, we mostly ignored specific regulatory processes such as involved in entering stationary phase, which may affect the quality of our predictions for poor growth media. Despite these limitations, our model recovers various types of data and empirical growth relations, highlighting the predictive power of our approach.

Our cell model links the stochasticity of intracellular mechanisms with growth variations observed in single cells and populations. Together with a novel theory to analyse stochastic cell and division dynamics, our work provides a framework to draw testable predictions and bring about a working understanding of the stochastic physiology in living cells.

## IV. METHODS

### a. List of reactions

The model consists of stochastic reactions, adapted from a previous deterministic model^22^, that represent transcription and degradation of mRNAs *m*_*y*_, their binding to free ribosomes *p*_*r*_ to form a ribosome-mRNA complex *c*_*y*_, and translation reactions synthesising a protein *p*_*y*_, where *y* ∈ {*t*, *e*, *r*, *q*}. To account for metabolism, we include uptake of an external nutrient *s* at fixed concentration by a transport protein (*p*_*t*_). The internalised nutrient *s*_int_ is then catabolised to produce resource molecules *a*. In summary, the stoichiometries and propensities of the reactions are:

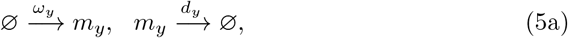

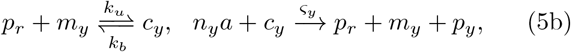

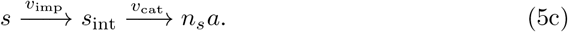

The propensities of mRNA degradation, ribosome binding and unbinding are modelled using mass action kinetics. Transcriptional and translational propensities depend on the resource *a* and follow 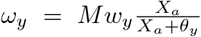 for *y* ∈ {*r*, *e*, *t*}, 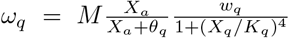 and 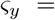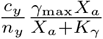 for *y* ∈ {*t*, *e*, *r*, *q*}, where *X* = *x*/*M* denote molecular concentrations. Nutrient uptake and metabolism is modelled using quasi-steady state kinetics via the propensities: *υ*_imp_ = *p*_*t*_*υ*_*t*_ and 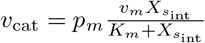.

The instantaneous growth rate can be obtained in closed form using Eq. (3) and is a product of the translation elongation rate and the concentration of translating ribosomes

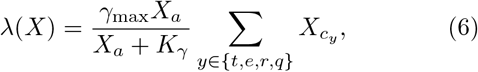

assuming mass is dominated by protein content. In the model, mean growth rate is varied through nutrient quality *n*_*s*_ describing the number of resource units produced per nutrient molecule.

To model operon architecture we replaced transcription, degradation, ribosome binding and translation reactions for transporters and enzyme species by the following set of reactions: 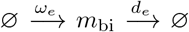, 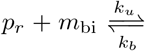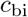, 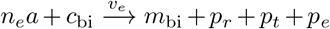 where *m*_bi_ is the bi-cistronic mRNA species coding for both transporter and enzyme proteins, *p*_*t*_ and *p*_*e*_, respectively.

Chloramphenicol effectively reduces the pool of elongating ribosomes by binding to ribosomes and preventing elongation^55^. We model this effect using an additional reaction:

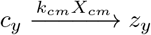

describing ribosome inhibition by the imposed drug concentration *X*_*cm*_ via a complex *z*_*y*_ that is no longer available to translation.

### b. Small noise approximation

Neglecting the noise terms in Eq. (4) reduces the Langevin equations to ODEs. In steady state the deterministic concentrations 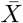 are characterised by the equations

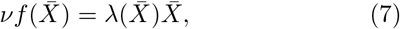

describing the balance between biomolecule synthesis by intracellular reactions and dilution of concentrations due to cell growth. In the limit of small fluctuations, i.e. the limit of large *M*, these concentrations describe the mean behaviour from which also the mean growth rate 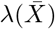 can be determined. Similarly, the average cell mass *M* increases exponentially between divisions with deterministic time-intervals 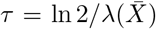. The average mass at cell birth follows from the delayed effect of initiation. Denoting the concentration of origins at which initiation of DNA-replication is triggered by *O*_*c*_, and the time required to complete replication and trigger cell division by *τ*_*C*+*D*_ (Fig. 1), the mean cell mass at birth and the number of origins are exponential functions of the mean growth rate

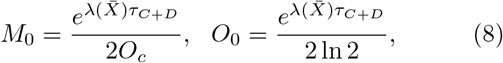

consistent with Donachie’s result^24^. This implies that the unit size (mass per number of origins) is constant and equal to ln2/*O*_*c*_. To compare against bulk data, we used the relation *M*_*bulk*_ = 2*M*_0_ ln 2 (neglecting size variation before and after division^56^).

The small noise approximation allows also to compute the size of growth fluctuations. The time-averaged concentration covariance 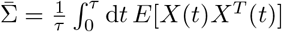 satisfies the following linear set of equations

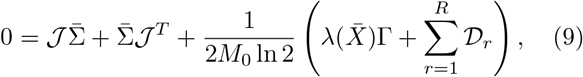

where 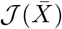 is the Jacobian of the deterministic ODEs and 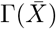 and 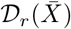 are the noise strengths of cell divisions and of the biochemical reactions, respectively (see SI Sec. IIB-D for details). These equations determine the size of fluctuations in growth rate via

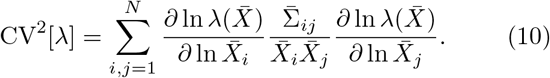

Following the lines of *Komorowski et al.*^39^, we analyse the sources of growth variations by decomposing Eq. (9) and (10) into contributions of cell divisions or groups of reactions (SI Sec. IIE).

### c. Model parametrisation

We parametrised the model with literature values for *E. coli* (SI Tab. I). For predicting the variability of single cells it is important to estimate the relative transcription rates of the involved mRNA species. To this end, we fitted the dependence of ribosomal mass fraction on mean growth rate against data from bulk experiments^8^ and the dependence of the CV^2^ [λ] on the mean against two recently published data sets of single-cell time-lapse microscopy^1,3^, allowing us to characterise a broad range of growth rates. We further constrained the maximal growth rate to 3.75 doublings per hour, equivalent to a minimal doubling time of 16 minutes. For the parameter estimation we used the small noise approximation of distribution moments combined with MCMC parameter sampling (SI Fig. S1, SI Sec. III).

### d. Stochastic simulations

We use a hybrid scheme that simulates reactions either using the next-reaction method or ODEs as described in^57^. To account for non-exponential reaction-time distributions^58^, we update propensities every 0.05 minutes. Supported by the predictions using the small noise approximation, it was sufficient to simulate only those reactions stochastically that change the lowly abundant mRNAs of transporters and enzymes and their corresponding ribosomal complexes. We determine growth rate using Eq. (3), which gives consistent results when measured at birth or division.

## AUTHOR CONTRIBUTIONS

PT, VD, AYW designed research; PT, GT, VD, AYW performed research; PT, AYW analysed data and wrote the paper.

## ACKNOWLEDGEMENTS

We thank Mauricio Barahona, Meriem El Karoui, Diego Oyarzún and Peter Swain for valuable feedback. PT gratefully acknowledges support from The Royal Commission for the Exhibition of 1851, and VD, AYW and GT from the ERC’s RULE project.

